# Pyrumthecina *Infraorder Nov.:* Revisiting the morphological diversity and identifying the phylogenetic home of *Argynnia* within Arcellinida (Tubulinea:Amoebozoa)

**DOI:** 10.1101/2025.08.20.671167

**Authors:** Alfredo L. Porfirio-Sousa, Maria Beatriz Gomes E Souza, Robert E. Jones, Matthew W. Brown, Daniel J. G. Lahr, Alexander K. Tice

## Abstract

Arcellinida is a diverse lineage of testate amoebae within Amoebozoa, whose evolutionary history has been clarified through phylogenomics. These efforts have led to a stable classification of the group into suborders and infraorders. However, several taxa, such as the genus *Argynnia*, remain unplaced due to ambiguous morphology and unresolved positions in single-gene phylogenies. In this study, we explore the diversity of *Argynnia* by presenting new records of species sampled across Brazil. To address its longstanding taxonomic uncertainty, we performed a phylogenomic analysis incorporating transcriptomic data from the Protist 10,000 Genomes isolate P10K-MW-000941 (P10K941). Through single-gene phylogenies, this isolate was shown to be closely related to *Argynnia*, which previously lacked genomic-level data. Here, we further confirm this relationship and present a 227-gene phylogenomic reconstruction that identifies the placement of P10K941 within Arcellinida, thereby resolving the phylogenetic placement of *Argynnia*. We obtained a curated, contaminant-free phylogenomic dataset for Arcellinida, including the P10K941, through the PhyloFisher workflow. Our analysis recovers P10K941 as a well-supported long-branch, sister to the infraorder Longithecina. Given this phylogenomic placement, along with *Argynnia*’s characteristic morphology and the consistent results of single-marker analyses placing it outside existing infraorders, we propose the Pyrumthecina infraorder *novum* and the family Argynniidae family *novum* to accommodate the P10K941 isolate and all *Argynnia* species. These findings resolve the phylogenetic position of *Argynnia* and allow for interpreting shell evolution and deep diversification patterns within Arcellinida. The recognition of Pyrumthecina also highlights the likely existence of deeply branching lineages among arcellinid taxa currently classified as *incertae sedis*.

## 1 Introduction

Phylogenomics has profoundly advanced our understanding of the evolution and diversification of Arcellinida, a diverse and ancient order of testate (shelled) amoebae within Amoebozoa (Lahr et al. 2019; Porfirio-Sousa et al. 2024). Recognized by their conspicuous shells, Arcellinida’s deep evolutionary relationships were first illuminated through the generation of whole-culture and single-cell transcriptomes, which enabled the reconstruction of a robust phylogenetic backbone (Lahr et al. 2019; Porter and Riedman 2019). Based on analyses of hundreds of genes across a wide taxonomic and evolutionary range, Arcellinida were classified into three suborders, Glutinoconcha, Organoconcha, and Phryganellina, and five infraorders within Glutinoconcha: Sphaerothecina, Longithecina, Excentrostoma, Hyalospheniformes, and Volnustoma (Lahr et al. 2019). Subsequent studies have corroborated the stability of this backbone and allowed for the identification of a novel infraorder (Cylindrothecina) and confident placement of previously unclassified taxa (González-Miguéns et al. 2022; Porfirio-Sousa et al. 2024). Together, these findings have significantly reshaped our understanding of Arcellinida and highlighted their value for exploring the deep evolutionary history of eukaryotes (Lahr et al. 2019; Porter and Riedman 2019; Lahr 2021; Porfirio-Sousa et al. 2024). Despite this significant progress, several arcellinids remain classified as *incertae sedis* (i.e., of uncertain placement), lacking a definitive position within the broader Arcellinida phylogeny (Kosakyan et al. 2016; Lahr et al. 2019).

Among the unresolved questions within Arcellinida is the genus *Argynnia* Vucetich, 1974, which stands out as a longstanding puzzle due to its ambiguous shell morphology and the consistent failure of single-gene phylogenies to place it within any well-established group (Vucetich 1974; Kosakyan et al. 2025). Characteristic of freshwater, aquatic plant rhizosphere, and moss habitats, *Argynnia* possesses a pear-shaped or ovoid, laterally compressed shell composed of mineral particles, often mixed with euglyphid plates or diatom fragments (Vucetich 1974; Meisterfeld 2002; Kosakyan et al. 2025). Originally classified within *Nebela* (Hyalospheniformes), *Argynnia* was later separated into its own genus due to its distinct morphology (Vucetich 1974; Meisterfeld 2002; Kosakyan et al. 2025). Besides their similarity to *Nebela* spp. (Hyalospheniformes), *Argynnia* also exhibits traits typical of *Difflugia* (Sphaerothecina), such as the presence of a structured organic cement and absence of a neck in the shell of most species (Meisterfeld 2002). This intermediate morphology has complicated the taxonomic placement of *Argynnia* since its early descriptions (Vucetich 1974; Meisterfeld 2002; Kosakyan et al. 2025). Reinforcing this ambiguity, single-gene phylogenies have consistently failed to resolve *Argynnia* within Hyalospheniformes, instead placing it outside any recognized infraorder (Gomaa et al. 2012; Kosakyan et al. 2012; González-Miguéns et al. 2022; Ribeiro et al. 2023). As a result, *Argynnia* remains assigned to Arcellinida *incertae sedis*, alongside several other Arcellinida taxa (Kosakyan et al. 2016; Lahr et al. 2019; Kosakyan et al. 2025). Despite its importance for understanding morphological evolution and diversification patterns within Arcellinida, owing to its distinct morphology and unresolved phylogenetic position, *Argynnia* remains poorly documented in some regions, and no genomic-level data had been available until recently. Specifically, although more than 20 species have been described worldwide, including from diverse regions of the Southern Hemisphere, only a few have been recorded in the literature to date in Brazil, the largest country in the Southern Hemisphere (Lansac-Tôha et al. 2001; Kosakyan et al. 2025).

Here, we document new records of multiple *Argynnia* species from an extensive Arcellinida sampling effort across Brazil and investigate the genus’s phylogenetic placement using newly available transcriptomic data. To resolve the longstanding phylogenetic uncertainty surrounding *Argynnia*, we conducted a 227-gene phylogenomic analysis of Arcellinida that includes the Protist 10,000 Genomes (P10K) isolate P10K-MW-000941 (hereafter P10K941). This pear-shaped taxon was originally collected from moss and identified as Hyalospheniidae sp. 3 P10K. Although lacking photo-documentation, P10K941 was recently identified as an *Argynnia* based on single-marker phylogenies (Porfirio-Sousa et al. 2025). In the present study, we performed BLAST similarity searches and phylogenetic analyses of SSU and COI sequences retrieved from the P10K941 transcriptomic data, confirming its close relationship to previously sequenced *Argynnia* specimens. Phylogenomic analysis using transcriptomic data revealed that P10K941 forms a well-supported, long-branch clade outside from all currently recognized Arcellinida infraorders, recovered as sister to Longithecina. To accommodate this novel lineage, we propose the new infraorder Pyrumthecina infraorder *nov*. and the new family Argynniidae fam. nov., comprising P10K941 and *Argynnia*. Ultimately, the establishment of Pyrumthecina not only provides a phylogenetic home for *Argynnia* but also points to the likely existence of deep evolutionary lineages hidden within the currently *incertae sedis* Arcellinida.

## 2 Methods

### 2.1 Sampling and microscopical observation of Argynnia specimens

We compiled records of *Argynnia* based on extensive sampling expeditions conducted across Brazil since 2012. These efforts covered seven Brazilian states and included the rhizosphere of aquatic plants, mosses, and leaf litter samples (**Supplementary Table 1**). In the laboratory, the vegetal material, comprising aquatic plant rhizospheres, mosses, and leaf litter, was washed with spring water to release testate amoebae. The washed material was then filtered using a 20 µm mesh to concentrate the specimens before microscopic screening. Concentrated samples were mounted on slides and examined under a Nikon Eclipse E200 light microscope equipped with a Prime Cam Intervision 12 MP camera system, used for photo-documenting the *Argynnia* specimens. The **Supplementary Table 1** provides detailed information on the sampling locations and conditions for each *Argynnia* specimen included in this study. The specimens presented here are part of a broader, long-term effort to document testate amoebae diversity in Brazil, led by Maria Beatriz Gomes e Souza. This ongoing initiative is showcased on the website https://tecamebas.com.br/, where additional images and records of *Argynnia* and other testate amoebae taxa can be accessed, with documentation dating back to the 1980s.

### 2.2 SSU and COI Phylogenetic and Similarity Analyses

We compiled small subunit ribosomal DNA/RNA (SSU rDNA/rRNA) and cytochrome c oxidase subunit I (COI) datasets representing a broad sampling of Arcellinida. These datasets were derived from previously constructed and curated dataset aimed at identifying the phylogenetic placement of Amoebozoan genomic data from the P10K project (Porfirio-Sousa et al. 2025). The original dataset included both SSU and COI sequences previously generated by several studies and sequences retrieved from genomic data of Amoebozoa available in the P10K database (Ribeiro et al. 2023; Porfirio-Sousa et al. 2024) (Porfirio-Sousa et al. 2025). From the datasets compiled in the present study, we performed phylogenetic reconstructions based on multiple sequence alignments (MSAs) of SSU and COI sequences, which were generated using MAFFT v7.490 with the E-INS-i algorithm and 1,000 refinement iterations (command: *mafft --genafpair --maxiterate 1000 input*.*fasta > output_aligned*.*fasta*) (Katoh and Standley 2013). Automated alignment trimming was performed with trimAl v1.5 using a gap threshold of 0.3 for SSU and 0.5 for COI (command: *trimal -in input_aligned*.*fasta -out output_aligned_trimmed*.*fasta -keepheader -gt [gap threshold]* (Capella-Gutiérrez et al. 2009). Phylogenetic trees were inferred from the trimmed alignments using the maximum likelihood method implemented in IQ-TREE v2.3.6. Model selection was performed using ModelFinder, and node support was assessed with 1,000 ultrafast bootstrap replicates and 1,000 SH-aLRT tests (command: *iqtree2 -s aligned_trimmed*.*fasta -alrt 1000 -bb 1000 -m TEST*) (Kalyaanamoorthy et al. 2017; Minh et al. 2020).

For the similarity assessment, we used the NCBI BLAST tool (Altschul et al. 1990) to compare the SSU and COI sequences of the P10K941 isolate against Arcellinida sequences in the NCBI nucleotide database. For SSU, we retrieved the top hit: *Argynnia dentistoma*. For COI, we retrieved the top hit: *Argynnia* sp. isolate Q6.

### 2.3 Phylogenomic Dataset Construction

We constructed our Arcellinida phylogenomic dataset using the database and tools provided by PhyloFisher v1.2.11 (Tice et al. 2021) following the detailed workflow available at https://thebrownlab.github.io/phylofisher-pages/detailed-example-workflow and the PhyloFisher protocol (Jones et al. 2024). Using the phylogenetically aware mode of *fisher*.*py*, we searched putative homologs of 240 target genes from the P10K941 isolate proteome available in the P10K database. As queries, we used orthologs previously identified in the PhyloFisher database from the tubulinid amoebozoans *Arcella uspiensis* (*intermedia*), *Cryptodifflugia operculata*, and *Copromyxa protea*, which were employed in HMMER and BLAST searches executed by *fisher*.*py* (Camacho et al. 2009; Mistry et al. 2013). For each of the 240 putative orthologs, sequences recovered by BLAST were incorporated into their respective alignments using *working_dataset_constructor*.*py*. These alignments already included orthologs and paralogs from diverse eukaryotic taxa included in the database provided by PhyloFisher v1.2.11 (Tice et al. 2021) and a comprehensive sampling of Arcellinida taxa from previous phylogenomic studies (Kang et al. 2017; Lahr et al. 2019; Porfirio-Sousa et al. 2024). Homolog trees were inferred from the extended alignments using *sgt_constructor*.*py*, and each tree was manually inspected in ParaSorter to confirm correct ortholog placement of P10K941 and to exclude any potential contaminant sequences from non-target eukaryotes. Final ortholog/paralog designations were applied to the PhyloFisher database using *apply_to_db*.*py*. To mitigate the effects of missing data in downstream analyses, we retained only orthologs present in at least 33% of the final taxon set. The resulting dataset was assembled using *prep_final_dataset*.*py* and *matrix_constructor*.*py* with default parameters. The final concatenated matrix used for phylogenetic analyses included 227 genes (68,836 amino acid sites) and was composed of 35 Arcellinida taxa alongside 13 closely related representatives from the major amoebozoan group Tubulinea (**Supplementary Table 2**).

### 2.4 Phylogenomic Analyses

We performed maximum likelihood (ML) phylogenetic reconstruction using our final matrix with IQ-TREE version 3.0.1 (Wong et al. 2025). We first inferred a tree under the ELM+C20+F+G model. This tree was then used as a guide to infer a Posterior Mean Site Frequency (PMSF) model using the ELM+C60+G+PMSF model in IQ-TREE. We assessed topological support for the resulting tree with 100 nonparametric bootstrap replicates under the PMSF model in IQ-TREE version 3.0.1 (Wong et al. 2025).

## 3 Results

### 3.1 Argynnia morphological and habitat diversity

Here we present seven species of *Argynnia* derived from the sampling effort of arcellinids across Brazil (**Fig. 1 and Supplementary Table 1**). These species exhibit a wide range of morphologies and sizes, including pear-shaped ovoid shells (**Fig. 1A–D**), more elongated shells lacking a neck (**Fig. 1E–F**), and shells with a distinct neck (**Fig. 1G**), as well as ovoid forms bearing spines (horns) **(Fig. 1H-J)**. Each morphotype displays a characteristic terminal shell aperture (pseudostome). Specimens of *Argynnia* have been collected from the rhizosphere of aquatic plants, mosses, and plant litter across diverse Brazilian regions and biomes. Sampling sites span seven states distributed among the South, Southeast, Central-West, and North regions of Brazil (**Fig. 1 and Supplementary Table 1**), encompassing three major biomes: the Atlantic Forest and Amazon (Tropical and Subtropical Moist Forests) and the Cerrado (Savanna).

**Fig. 1.**
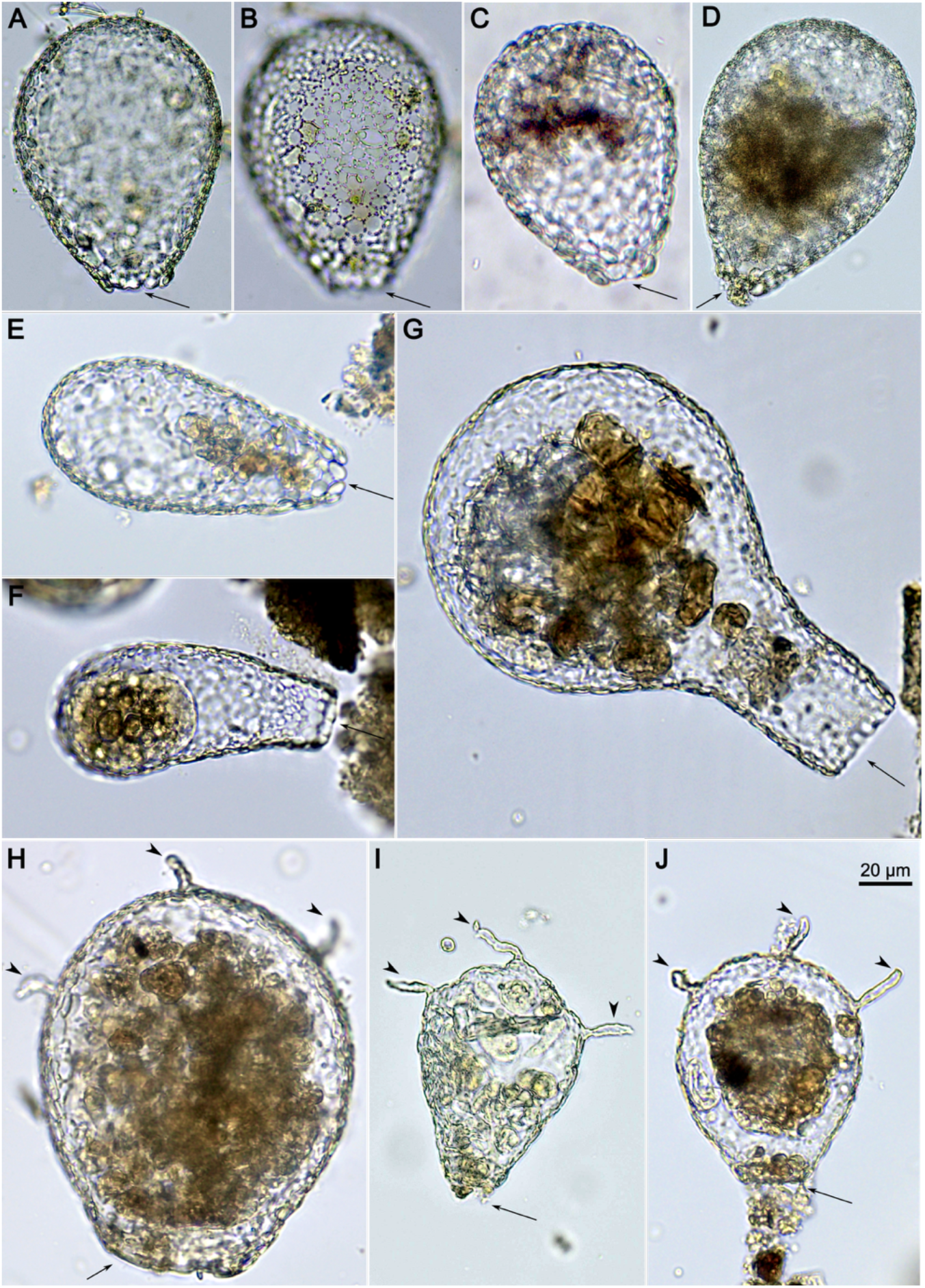
Representative specimens of *Argynnia species* collected in Brazil. *Argynnia* is a genus recognized by a pear-shape or ovoid shell, usually compressed, covered with scale plates of different shapes and exogenous particle, including euglyphid plates or diatom fragments. **A-C**. *Argynnia dentistoma* (Penard, 1890) Vucetich, 1974. Ovoid shell with an oval pseudostome (shell aperture) composed of plates forming a denticulated (small toothlike protrusions) boarder. **D**. *Argynnia dentistoma* var. *lacustris* Wailes, 1919. It differs from *A. dentistoma* by having a broader and more circular shell. **E**. *Argynnia oblonga* Gauthier-Lièvre, 1953. Shell ovoid-elongated, with a pseudostome composed of plates forming a denticulated margin, lacking a defined collar around the aperture. **F**. *Argynnia repanda* Jung, 1942. Shell elongated and slightly curved, with an oval pseudostome composed of larger plates. **G**. *Argynnia gertrudeana* Jung, 1942. Shell pyriform, bottle-shaped, with a neck of variable length and width, and an oval pseudostome. **H**. *Argynnia spicata* Wailes, 1913. Shell broadly ovoid, sometimes almost circular, with a series of 2 to 8 horns of variable sizes on the shell surface. Aperture broad and oval. Differs from *Argynnia caudata* by its larger size and the more circular shape of the shell. **I**. *Argynnia kundulungui* Van Oye, 1959. Shell ovoid or pyriform, with a series of 2 to 4 long horns arranged equidistantly. Oval pseudostome reinforced with distinct plates. Differs from *A. caudata* by the shape of the shell, the arrangement of the horns, and the distinct characteristics of the pseudostome. **J**. *Argynnia caudata* Leidy, 1879. Shell broadly pyriform, sometimes almost circular, with a series of 2 to 6 horns of varying sizes along the shell margin. The pseudostome is circular and crenulated with a thin lip. The shell characteristics described for each species represent the features observed and used for species identification; not all characteristics are visible in the representative picture. Arrows show the pseudostome region; arrowheads show the shell’s horns. The images are to scale, a 20 μm scale bar is shown.

### 3.2 SSU and COI phylogenetics and similarity assessment

The inferred SSU maximum likelihood (ML) tree comprises 57 taxa, representing a broad sampling of Arcellinida diversity, with three Euamoebida taxa included as outgroup (**Fig. 2B**). In this tree, Arcellinida infraorders are recovered with high support (SH-aLRT ≥ 99.6 /UFBoot ≥ 100), with exception of the Organoconcha infraorder that is recovered with a lower UFBoot (SH-aLRT = 91.6 /UFBoot = 81) (**Fig. 2B**). This recovery of Arcellinida infraorders allows for the robust placement of most SSU sequences from the P10K dataset within these clades, except for the P10K941 isolate (**Fig. 2B**). Specifically, the SSU sequence of the P10K941 isolate is recovered with high support (SH-aLRT ≥ 91.9 /UFBoot ≥ 97) as sister to Argynnia dentistoma (**Fig. 2B**). The clade comprising P10K941 and Argynnia dentistoma does not nest within any established infraorder and is recovered as sister to the combined clades Longithecina + Sphaerothecina (**Fig. 2B**).

**Fig. 2.**
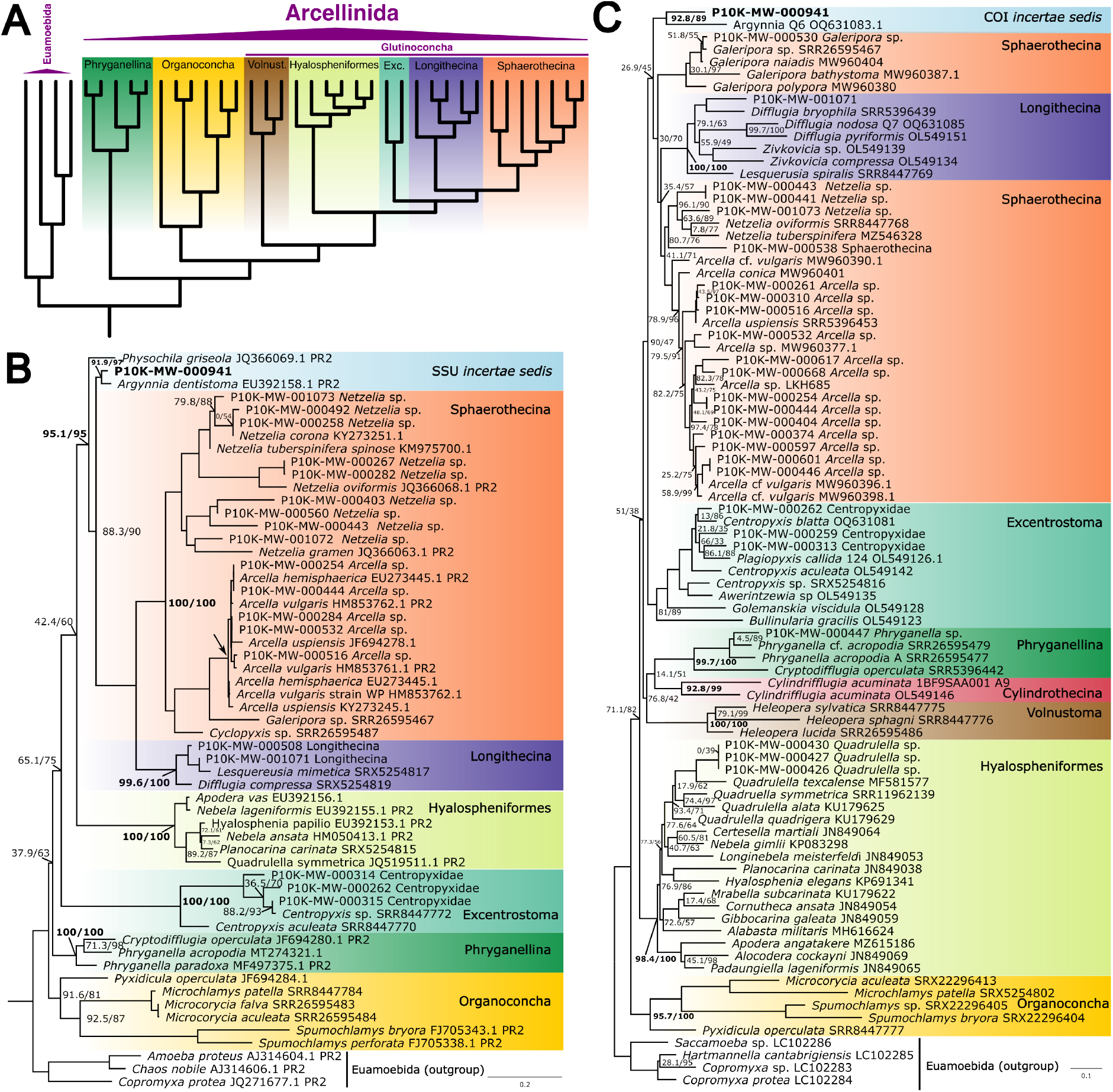
Diagrammatic tree of the order Arcellinida and single-marker phylogenetic trees placing the P10K941 isolate. **A**. Diagrammatic tree of the order Arcellinida based on the phylogenomic tree from Porfirio-Sousa et al., 2024. Abbreviations: Vonust., Volnustoma; Exc., Excentrotoma. **B**. Phylogenetic tree using SSU, a more conserved marker, showing phylogenetic affinity between the P10K941 isolate and *Argynnia dentistoma*. **C**. Phylogenetic tree using COI, a more variable mitochondrial marker, showing phylogenetic affinity between the P10K941 isolate and *Argynnia* sp. Support values are reported as SH-aLRT /UFBoot, with values ≥80/95 considered indicative of strong support. For clarity, high-support values are omitted in this figure, except for key nodes related to *Argynnia* or those delimiting the infraorders, which are shown in bold. Support values are also omitted for nodes above the one indicated by an arrow, which are mostly represented by flat branches. The complete tree, including all support values, is shown in Supplementary Figs. 1 and 2.

The inferred COI maximum likelihood (ML) tree comprises 87 taxa, also representing a broad sampling of Arcellinida diversity, with four Euamoebida taxa included as outgroup (**Fig. 2C )**. In this tree, most Arcellinida infraorders are recovered with high support (SH-aLRT ≥ 92.8 /UFBoot ≥ 99), except for Excentrostoma, which is recovered with high SH-aLRT and nearly high UFBoot support, and Sphaerothecina, which appears split by the Longithecina infraorder **(Fig. 2C)**. The COI tree allows for the robust placement of most COI sequences from the P10K dataset within the sampled infraorders. Specifically, the COI sequence of the P10K941 isolate is recovered with high SH-aLRT and nearly high UFBoot support (SH-aLRT ≥ 92.8 /UFBoot ≥ 99) as sister to *Argynnia* sp. isolate Q6. The clade comprising P10K941 and *Argynnia* sp. Q6 does not nest within any established infraorder and is recovered with low support as sister to the combined clades Longithecina + Sphaerothecina (**Fig. 2C**).

The similarity assessment based on the SSU sequence of the P10K941 isolate identifies *Argynnia dentistoma* (GenBank accession EU392158.1) as the top match, with an alignment of 1,033 base pairs showing 96% identity and 0.1% gaps (**Supplementary Table 3**). The similarity search based on the COI sequence of P10K941 identifies *Argynnia* sp. Q6 (GenBank accession OQ631083.1) as the top match, with an alignment of 567 base pairs showing 83% identity and 1% gaps (**Supplementary Table 4**).

### 3.3 Phylogenomic reconstruction

The inferred phylogenomic tree, comprising 48 taxa and 227 genes (68,836 amino acid sites), fully resolves Arcellinida and recovers with full support its three suborders (Phryganellina, Organoconcha, and Glutinoconcha), as well as the five infraorders within Glutinoconcha (**Fig. 3**). The P10K941 isolate does not nest within any established infraorder and is recovered with nearly full support as a long-branch sister to the Longithecina infraorder within Glutinoconcha (**Fig. 3**).

**Fig. 3.**
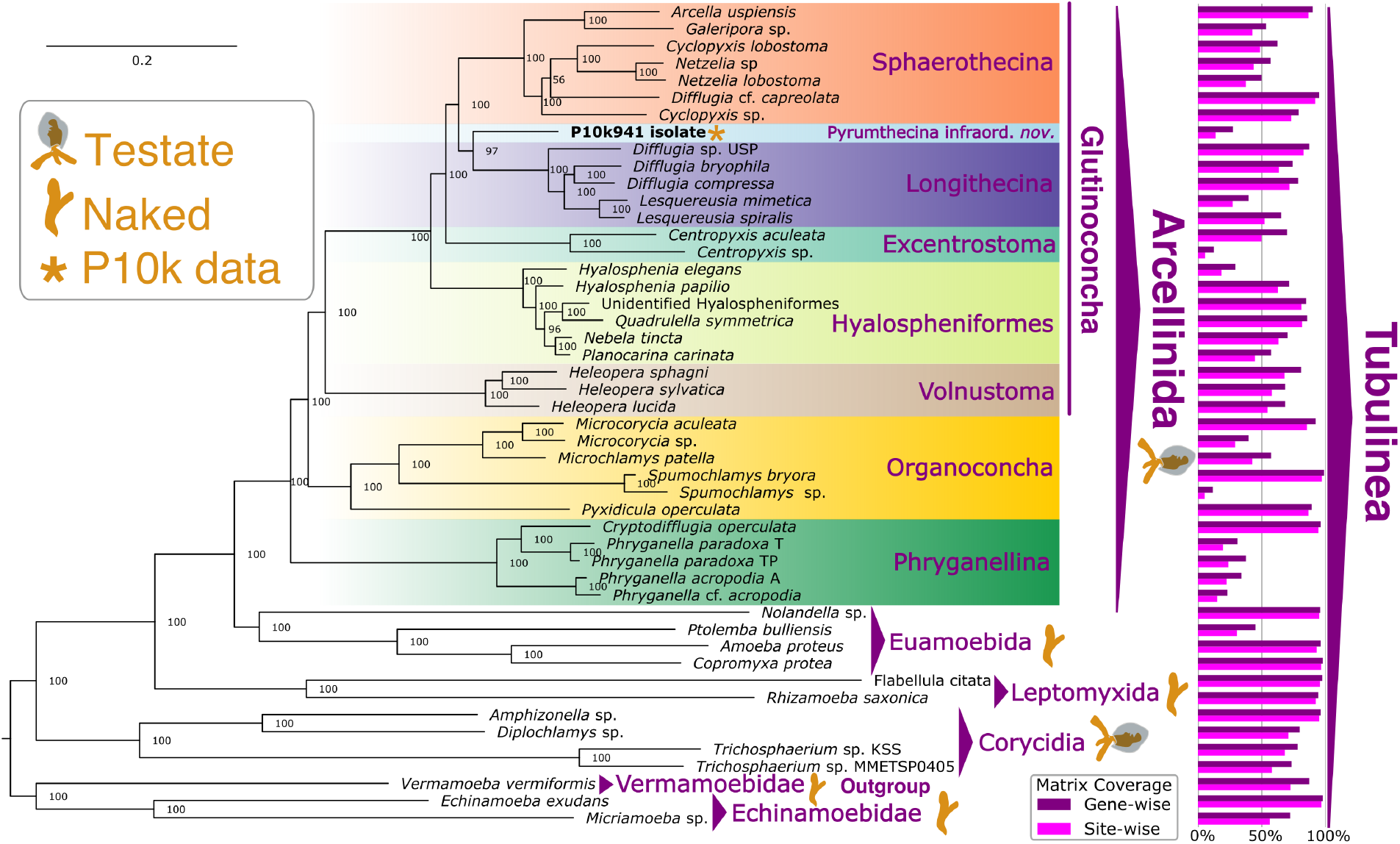
Phylogenomic tree of Tubulinea. Phylogeny of Arcellinida and closely related Tubulinea based on 227 genes (68,836 amino acid sites). The tree was initially built using IQ-TREE v. 3.0.1 under the ELM+C20+G4 model of protein evolution and further used to infer a posterior means site frequency model using the ML model ELM+C60+G4+PMSF. Topological support was assessed by 100 MLRB replicates. The bar plots on the right side of the figure show the percentage completeness of the dataset available for each taxon, based on the total number of genes and sites in our phylogenomic tree matrix.

## 4 Discussion

### 4.1 Argynnia diversity and distribution

The *Argynnia* species reported in this study expand the known distribution of the genus in Brazil and highlight its remarkable morphological and ecological diversity. Until recently, records in the literature of *Argynnia* in Brazil were limited to a few species, including *A. caudata, A. dentistoma*, and *A. vitre*a (Lansac-Tôha et al. 2001). Globally, although some species exhibit cosmopolitan or hemispheric distributions, appearing in both the Northern and Southern Hemispheres, many *Argynnia* species appear to be restricted to the Southern Hemisphere, particularly in South America (Vucetich 1974; Meisterfeld 2002; Heger et al. 2009; Kosakyan et al. 2025). Within Brazil, the presence of *Argynnia* across diverse geographic regions, comprising three major biomes, underscores the ecological range of this genus. This ecological breadth is matched by considerable morphological diversity, as evidenced by the variety of distinctive shell shapes represented by the genus. Such patterns of ecological and morphological variation are consistent with observations from other genera within Arcellinida, especially those that have undergone taxonomic revisions based on detailed morphological and molecular analyses (Kosakyan et al. 2016, 2025; Dumack et al. 2019, 2020; González-Miguéns et al. 2022). Specifically, *Argynnia* includes several species and species varieties that require further detailed morphological and molecular investigation. For instance, *Argynnia antarctica* has traditionally been considered an example of a testate amoeba with a restricted geographical distribution, originally described only from Marion Island in the southern Indian Ocean (Grospietsch 1971; Heger et al. 2009). However, more extensive morphological and morphometric analyses have shown that the morphometric range of the cosmopolitan species *Argynnia dentistoma* overlaps with that reported for *A. antarctica*, suggesting that they may be synonyms and that *A. antarctica* does not represent an endemic taxon (Heger et al. 2009). Nevertheless, a more comprehensive survey of *A. antarctica* and *A. dentistoma*, including molecular data, is still lacking. Similarly, *A. dentistoma* is currently recognized in several varieties, some of which have restricted geographical distributions and are known from only a few records, thus also requiring further study (Playfair 1918; Vucetich 1974; Dura 1986; Kosakyan et al. 2025) (Playfair, 1917; Vucetich, 1973; Decloitre, 1977; Cerdá, 1986). Regarding molecular data, *Argynnia dentistoma* is the only recognized species in the genus with publicly available data, in addition to *Argynnia* sp. Q6 and the P10K941 isolate. Consequently, further morphological assessment and molecular sampling of *Argynnia* are necessary to bring novel understanding of the diversity and geographical distribution of this genus and for the Arcellinida group.

### 4.2 Phylogenetic placement of Argynnia

The congruent results from SSU and COI phylogenetic reconstruction and similarity assessment support a close taxonomic affinity between the P10K941 isolate and the genus *Argynnia*. Both markers consistently recover P10K941 and *Argynnia* as sister lineages, with SSU showing higher sequence identity and shorter branches, as expected for its slower substitution rate (Fig. 2 B-C). Despite robustly recovering the Arcellinida infraorders and precisely placing the arcellinid P10K sequences considered in our reconstructions, the SSU and COI trees fail to place *Argynnia* and P10K941 within any established group. Both SSU and COI place them as basal groups of the clade composed of Sphaerothecina and Longithecina. This pattern is consistent with numerous previous studies based on single-gene phylogenies with varying taxon sampling, which also recovered *Argynnia* outside recognized infraorders and basal to a clade including *Arcella, Netzelia*, and *Difflugia*, genera that currently compose Sphaerothecina and Longithecina infraorders (Lara et al. 2008; Gomaa et al. 2012; Kosakyan et al. 2012; González-Miguéns et al. 2022; Ribeiro et al. 2023).

Based on the congruency between our phylogenetic reconstructions and sequence similarity assessments, in line with previous studies, we propose that the P10K941 isolate likely represents a member of *Argynnia*, or more conservatively, at the very least a closely related yet uncharacterized genus. Although the P10K database does not provide a voucher image associated with this isolate, it describes P10K941 as a specimen with a pear-shaped shell collected from moss. These morphological and habitat characteristics are consistent with those of *Argynnia* and their phylogenetic affinity as inferred from molecular data (Vucetich 1974; Kosakyan et al. 2025; Porfirio-Sousa et al. 2025). Moreover, since SSU and COI markers have consistently proven effective at recovering Arcellinida infraorders when broad taxon sampling is applied (González-Miguéns et al. 2022; Ribeiro et al. 2023), their inability to place P10K941 and *Argynnia* within any established infraorder supports the hypothesis that these taxa may represent a distinct clade within Arcellinida. Given the availability of transcriptomic data for the P10K941 isolate, this interpretation could be further tested through phylogenomic analysis.

Our phylogenomic reconstruction based on 227 genes is fully congruent with previous phylogenomic studies and reinforces the current higher-level classification of Arcellinida (Lahr et al. 2019; Porfirio-Sousa et al. 2024). The recovery of the three recognized suborders (Phryganellina, Organoconcha, and Glutinoconcha) and all five infraorders within Glutinoconcha, with maximal support across all nodes, confirms that the overall backbone of Arcellinida is currently well resolved and stable. Within this framework, the P10K941 isolate is recovered as a long-branch lineage, with nearly full support as sister to the infraorder Longithecina. This placement is to some extent congruent with the SSU and COI phylogenies, which place P10K941 and *Argynnia* basally to the clade composed of Longithecina and Sphaerothecina. The recovery of P10K941 as a distinct lineage in the phylogenomic tree demonstrates that the inability of single-gene analyses to place P10K941 and *Argynnia* within a known infraorder is not solely due to insufficient phylogenetic signal in these markers, but rather reflects the evolutionary distinctiveness of these taxa.

Taken together, the consistent phylogenetic signal from SSU, COI, and phylogenomic analyses, alongside the unique combination of morphological features of *Argynnia* (Vucetich 1974; Kosakyan et al. 2025), justify their recognition as a lineage separate from any currently described infraorder. Since its original description, *Argynnia* has been considered a morphological mosaic, exhibiting traits associated with both *Nebela*-like and *Difflugia*-like taxa (Vucetich 1974; Kosakyan et al. 2025). Now, integrating robust molecular evidence, we interpret that Argynnia and P10K941 belong to a distinct lineage within Arcellinida. Their characteristic morphology is likely related to the retention of shell traits inherited from a common ancestor shared with the Longithecina infraorder, as well as the convergent evolution of shell traits observed in Hyalospheniformes, an infraorder more distantly related to the genus *Argynnia*. To accommodate this distinctiveness, we propose the establishment of Pyrumthecina infraorder *novum* and Argynniidae Family *novum*, which includes P10K941 and *Argynnia*. It is worth noting that Pyrumthecina and Argynniidae are currently established phylogenomically based on a single isolate. Since their monophyly cannot be formally tested with fewer than two terminals, the inclusion of additional taxa within *Argynnia* will be necessary to further investigate this new infraorder and assess Argynniidae monophyly. Nonetheless, the combined evidence from morphology, SSU, COI, and the long-branch phylogenomic placement, consistent with patterns observed in other well-established infraorders, supports the erection of Pyrumthecina and provides a clear target for future sampling, allowing the testing of its monophyly and the refinement of the Arcellinida tree of life.

### 4.3 Taxonomic actions

Here we continue to reconcile the prevailing usage of the PhyloCode in higher-level protistan classifications and the traditional classification of testate amoebae regulated by the International Code of Zoological Nomenclature, as previously established for Arcellinida (Lahr et al. 2019). Accordingly, we adopt the hierarchical system proposed by Adl and collaborators (Adl et al. 2012) for indicating levels of inclusiveness among protist groups, in which increasing numbers of bullet points correspond to increasingly nested and less inclusive ranks. We also suggest how these hierarchical levels align with traditional Linnaean ranks: the highest taxon treated here (Tubulinea) is indicated by “•” and is at the Linnean rank of “Class”, Arcellinida is indicated by “•••” and is at the level of “Order”, those indicated by “••••” are at the level of “Suborder”, those indicated by “•••••” are at the level of “Infraorder”, and those indicated by “••••••” are at the level of “Family”.

Amorphea Adl et al. 2012

Amoebozoa Lühe 1913, sensu Cavalier-Smith 1998

•Class Tubulinea Smirnov et al. 2005

•• Clade Elardia Kang et al. 2017

••• Order Arcellinida Kent 1880

•••• Suborder Glutinoconcha Lahr et al. 2019

••••• Infraorder Pyrumthecina infraord. nov. Porfirio-Sousa, Gomes E Souza, Jones, Brown, Lahr, and Tice 2025

•••••• Family Argynniidae *fam. nov*. Porfirio-Sousa, Gomes E Souza, Jones, Brown, Lahr, and Tice 2025

*Node-based definition:* the clade originating with the first ancestor of *Argynnia* and isolate P10K941 that is not the ancestor of Longithecina.

*Diagnosis:* can be diagnosed by its specific sequences of the nuclear and mitochondrial DNA markers (SSU and COI) and by its phylogenetic placement.

*Type Family: Argynniidae fam. nov*. Porfirio-Sousa, Gomes E Souza, Jones, Brown, Lahr, and Tice 2025

#### Remarks

The genus *Argynnia* Vucetich 1974 has remained a taxonomic enigma within Arcellinida due to their distinct shell morphology and lack of phylogenetic resolution based on single-marker reconstructions. Originally classified in the family Nebelidae because of its pear-shaped, laterally compressed test composed of mineral particles and euglyphid plates, features resembling *Nebela, Argynnia* genus establishment was initially invalid due to the lack of a type species. This was later corrected with the designation of *Argynnia schwabei* as the type (Vucetich 1974; Meisterfeld 2002; Kosakyan et al. 2025). Despite morphological similarities to nebelid genera, *Argynnia* also exhibits traits typical of *Difflugia* (Sphaerothecina) and molecular phylogenies based on SSU and COI have consistently failed to place *Argynnia* within Hyalospheniformes or any recognized infraorder. Consequently, *Argynnia*’s evolutionary relationships remained unresolved. The close relationship between *Argynnia* and the P10K941 isolate, combined with the phylogenomic placement of the P10K941 isolate, illuminate the enigma and clarifies why single-gene markers have failed to place *Argynnia*, they comprise an independent deeper branch within Arcellinida, accommodated by the erection of Pyrumthecina infraord. nov. as a sister infraorder to Longithecina.

#### Etymology

Lt. pyrum – pear (fruit), Lt. theca - shell.

*Included Taxa:* Family Argynniidae Fam. Nov., Genera *Argynnia*.

## 1 Conclusions

This study expands the identification and photo-documentation of *Argynnia* available in the literature and elucidates the longstanding phylogenetic uncertainty surrounding this genus by establishing its placement within Arcellinida. By reporting *Argynnia* species identified from Brazil, we highlight the morphological and ecological diversity associated with this genus. Using a dataset of 227 genes for a phylogenomic reconstruction of the Arcellinida tree, we recovered P10K941, an arcellinid isolate that is phylogenetically, morphologically, and ecologically consistent with *Argynnia*, as a long-branch lineage sister to Longithecina within Glutinoconcha. This phylogenomic reconstruction is congruent with phylogenetic analyses based on SSU and COI markers, which consistently place both P10K941 and previously sequenced *Argynnia* as a lineage outside any previously described arcellinid infraorder. This congruence across molecular datasets, combined with the morphological distinctiveness of *Argynnia*, supports the recognition of a new infraorder within Glutinoconcha: Pyrumthecina *infraord. nov*. The establishment of Pyrumthecina resolves the uncertain taxonomic status of the currently described *Argynnia* species and the P10K941 isolate within Arcellinida. More broadly, the establishment of Pyrumthecina underscores the potential for uncovering additional major clades within Arcellinida. As genomic data become available for more morphologically diverse testate amoebae, other unplaced or enigmatic taxa are likely to be recognized as distinct evolutionary lineages. This continued expansion of taxon-rich phylogenomic frameworks will be essential to refining the evolutionary history of Arcellinida and broadening our understanding not only of testate amoeba evolution but of eukaryotic evolution as a whole.

## Availability of data and materials

All compiled and curated SSU, COI, and phylogenomic datasets, including single-protein untrimmed and trimmed alignments for each of the 227 homologous genes used in constructing our phylogenomic matrix, are publicly available on FigShare at https://doi.org/10.6084/m9.figshare.29944340

## Competing interests

The authors declare that they have no known competing interest.

## CRediT authorship contribution statement

**Alfredo L. Porfirio-Sousa:** Conceptualization, Data curation, Visualization, Formal analysis, Investigation, Writing – original draft, Writing – review & editing. **Robert E. Jones**: Investigation, Data curation, Writing – review & editing. **Maria Beatriz Gomes E Souza:** Data curation, Sampling, Photo-documentation, Writing – review & editing. **Matthew W. Brown:** Investigation, Writing – review & editing. **Daniel Lahr**: Investigation, Writing – review & editing. **Alexander K. Tice:** Conceptualization, Data curation, Visualization, Formal analysis, Investigation, Writing – original draft, Writing – review & editing, Resources, Supervision, Funding acquisition.

## Declaration of generative AI in scientific writing

During the preparation of this work the authors used ChatGPT to improve the readability and language of the manuscript. After using this tool, the authors reviewed and edited the content as needed and take full responsibility for the content of the published article.

## Funding

A.L.P-S and R.E.J were supported by startup funds provided to A.K.T. by Texas Tech University. This work was supported by the National Science Foundation Division of Environmental Biology (2100888) awarded to M.W.B., D.J.G.L. is supported by a FAPESP award #2019/22815-2.

## Acknowledgments

We thank Ferry Siemensma for his valuable discussions and taxonomic suggestions during the preparation of this manuscript. The authors acknowledge the High Performance Computing Center (HPCC) at Texas Tech University for providing computational resources that have contributed to the research results reported within this paper. URL: http://www.hpcc.ttu.edu

## Notes

https://doi.org/10.6084/m9.figshare.29944340

### Competing Interest Statement

The authors have declared no competing interest.

## References

Adl S. M., Simpson A. G. B., Lane C. E., Lukeš J., Bass D., Bowser S. S., Brown M. W., Burki F., Dunthorn M., Hampl V., Heiss A., Hoppenrath M., Lara E., le Gall L., Lynn D. H., McManus H., Mitchell E. A. D., Mozley-Stanridge S. E., Parfrey L. W., Pawlowski J., Rueckert S., Shadwick L., Schoch C. L., Smirnov A. & Spiegel F. W. 2012. The Revised Classification of Eukaryotes. Journal of Eukaryotic Microbiology, 59:429–514.

Altschul S. F., Gish W., Miller W., Myers E. W. & Lipman D. J. 1990. Basic local alignment search tool. Journal of Molecular Biology, 215:403–410.

Camacho C., Coulouris G., Avagyan V., Ma N., Papadopoulos J., Bealer K. & Madden T. L. 2009. BLAST+: architecture and applications. BMC Bioinformatics, 10:421.

Capella-Gutiérrez S., Silla-Martínez J.M. & Gabaldón T. 2009. trimAl: a tool for automated alignment trimming in large-scale phylogenetic analyses. Bioinformatics, 25:1972–1973.

Dura F. J. C. 1986. Contribucion al estudio de los protozosa rhizopoda (amebas testaceas) de la cordillera andina. Universitat de Barcelona. Available from: https://dialnet.unirioja.es/servlet/tesis?codigo=245079

Dumack K., Görzen D., González-Miguéns R., Siemensma F., Lahr D. J. G., Lara E. & Bonkowski M. 2020. Molecular investigation of Phryganella acropodia Hertwig et Lesser, 1874 (Arcellinida, Amoebozoa). European Journal of Protistology, 75:125707.

Dumack K., Kahlich C., Lahr D.J.G. & Bonkowski M. 2019. Reinvestigation of Phryganella paradoxa (Arcellinida, Amoebozoa) Penard 1902. Journal of Eukaryotic Microbiology, 66:232–243.

Gomaa F., Todorov M., Heger T. J., Mitchell E.A.D. & Lara E. 2012. SSU rRNA Phylogeny of Arcellinida (Amoebozoa) Reveals that the Largest Arcellinid Genus, Difflugia Leclerc 1815, is not Monophyletic. Protist, 163:389–399.

González-Miguéns R., Todorov M., Blandenier Q., Duckert C., Porfirio-Sousa A. L., Ribeiro G. M., Ramos D., Lahr D. J. G., Buckley D. & Lara E. 2022. Deconstructing Difflugia: The tangled evolution of lobose testate amoebae shells (Amoebozoa: Arcellinida) illustrates the importance of convergent evolution in protist phylogeny. Molecular Phylogenetics and Evolution, 175:107557.

Grospietsch T. 1971. Rhizopoda – Beitrag zur Ökologie der testaceen Rhizopoden von Marion Island. Marion and Prince Edward Islands. :411–419.

Heger T. J., Mitchell E. a. D., Ledeganck P., Vincke S., Van de Vijver B. & Beyens L. 2009. The curse of taxonomic uncertainty in biogeographical studies of free-living terrestrial protists: a case study of testate amoebae from Amsterdam Island. Journal of Biogeography, 36:1551–1560.

Jones R. E., Tice A. K., Eliáš M., Eme L., Kolísko M., Nenarokov S., Pánek T., Rokas A., Salomaki E., Strassert J. F. H., Shen X.-X., Žihala D. & Brown M. W. 2024. Create, Analyze, and Visualize Phylogenomic Datasets Using PhyloFisher. Current Protocols, 4:e969.

Kalyaanamoorthy S., Minh B. Q., Wong T. K. F., von Haeseler A. & Jermiin L. S. 2017. ModelFinder: fast model selection for accurate phylogenetic estimates. Nat Methods, 14:587–589.

Kang S., Tice A. K., Spiegel F. W., Silberman J. D., Pánek T., Čepička I., Kostka M., Kosakyan A., Alcântara D. M. C., Roger A. J., Shadwick L. L., Smirnov A., Kudryavtsev A., Lahr D.J.G. & Brown M. W. 2017. Between a Pod and a Hard Test: The Deep Evolution of Amoebae. Molecular Biology and Evolution, 34:2258–2270.

Katoh K. & Standley D. M. 2013. MAFFT Multiple Sequence Alignment Software Version 7: Improvements in Performance and Usability. Mol Biol Evol, 30:772–780.

Kosakyan A., Gomaa F., Lara E. & Lahr D. J. G. 2016. Current and future perspectives on the systematics, taxonomy and nomenclature of testate amoebae. European Journal of Protistology, 55:105–117.

Kosakyan A., Heger T. J., Leander B. S., Todorov M., Mitchell E.A.D. & Lara E. 2012. COI Barcoding of Nebelid Testate Amoebae (Amoebozoa: Arcellinida): Extensive Cryptic Diversity and Redefinition of the Hyalospheniidae Schultze. Protist, 163:415–434.

Kosakyan A., Ralf M., Enrique L., Clément D. & Mitchell Edward A. D. 2025. A taxonomic monograph of hyalospheniid testate amoebae Amoebozoa: Arcellinida: Hyalospheniformes. CHE. Available from: https://iris.unimore.it/handle/11380/1368035

Lahr D. J. 2021. An emerging paradigm for the origin and evolution of shelled amoebae, integrating advances from molecular phylogenetics, morphology and paleontology. Mem. Inst. Oswaldo Cruz, 116:e200620.

Lahr D. J. G., Kosakyan A., Lara E., Mitchell E. A. D., Morais L., Porfirio-Sousa A. L., Ribeiro G. M., Tice A. K., Pánek T., Kang S. & Brown M. W. 2019. Phylogenomics and Morphological Reconstruction of Arcellinida Testate Amoebae Highlight Diversity of Microbial Eukaryotes in the Neoproterozoic. Current Biology, 29:991-1001.e3.

Lansac-Tôha F. A., Velho L. F. M., Takahashi E. M., Aoyagui AS.M. & Bonecker C. C. 2001. Ocorrência de tecamebas (Protozoa, Rhizopoda) em águas continentais brasileiras. V. Famílias Hyalospheniidae, Plagiopyxidae, Microcoryciidae, Cryptodifflugiidae, Phryganelidae, Euglyphidae, Trinematiidae e Cyphoderiidae. Acta Scientiarum. Biological Sciences, 23:333–347.

Lara E., Heger T. J., Ekelund F., Lamentowicz M. & Mitchell E. A. D. 2008. Ribosomal RNA Genes Challenge the Monophyly of the Hyalospheniidae (Amoebozoa: Arcellinida). Protist, 159:165–176.

Meisterfeld R. 2002. Arcellinida. In: An Illustrated Guide to the Protozoa. Vol. 2. Second. Society of Protozoologists, Lawrence, Kansas, U.S.A. p. 827–859.

Minh B. Q., Schmidt H. A., Chernomor O., Schrempf D., Woodhams M. D., von Haeseler A. & Lanfear R. 2020. IQ-TREE 2: New Models and Efficient Methods for Phylogenetic Inference in the Genomic Era. Mol Biol Evol, 37:1530–1534.

Mistry J., Finn R. D., Eddy S. R., Bateman A. & Punta M. 2013. Challenges in homology search: HMMER3 and convergent evolution of coiled-coil regions. Nucleic Acids Res, 41:e121.

Playfair G. I. 1918. Rhizopods of Sydney and Lismore. Proceedings of the Linnean Society of New South Wales, 42:633–675.

Porfirio-Sousa A. L., Jones R. E., Brown M.W. & Lahr D. J. G., Tice A. K. 2025. Phylogenetic placement and contamination screening of Amoebozoa genomic data from the Protist 10,000 Genomes (P10K) Database. bioRxiv.

Porfirio-Sousa A. L., Tice A. K., Morais L., Ribeiro G. M., Blandenier Q., Dumack K., Eglit Y., Fry N. W., Gomes E Souza M. B., Henderson T. C., Kleitz-Singleton F., Singer D., Brown M.W. & Lahr D. J. G. 2024. Amoebozoan testate amoebae illuminate the diversity of heterotrophs and the complexity of ecosystems throughout geological time. Proceedings of the National Academy of Sciences, 121:e2319628121.

Porter S.M. & Riedman L. A. 2019. Evolution: Ancient Fossilized Amoebae Find Their Home in the Tree. Current Biology, 29:R212–R215.

Ribeiro G. M., Useros F., Dumack K., González-Miguéns R., Siemensma F., Porfírio-Sousa A. L., Soler-Zamora C., Pedro Barbosa Alcino J., Lahr D.J.G. & Lara E. 2023. Expansion of the cytochrome C oxidase subunit I database and description of four new lobose testate amoebae species (Amoebozoa; Arcellinida). European Journal of Protistology, 91:126013.

Tice A. K., Žihala D., Pánek T., Jones R. E., Salomaki E. D., Nenarokov S., Burki F., Eliáš M., Eme L., Roger A. J., Rokas A., Shen X.-X., Strassert J. F. H., Kolísko M. & Brown M. W. 2021. PhyloFisher: A phylogenomic package for resolving eukaryotic relationships. PLOS Biology, 19:e3001365.

Vucetich M. C. 1974. Comentarios criticos sobre Argynnia Jung, 1942 (Rhizopoda, Testacea). Neotropica.

Wong T. K. F., Ly-Trong N., Ren H., Baños H., Roger A. J., Susko E., Bielow C., Maio N. D., Goldman N., Hahn M. W., Huttley G., Lanfear R. & Minh B. Q. 2025. IQ-TREE 3: Phylogenomic Inference Software using Complex Evolutionary Models. Available from: https://ecoevorxiv.org/repository/view/8916/

