## Supplementary Material for "Pyrumthecina *Infraorder Nov.:* Revisiting the morphological diversity and identifying the phylogenetic home of *Argynnia* within Arcellinida (Tubulinea:Amoebozoa)"

**This file includes:**

Supplementary Tables 1 – 4 legends

Supplementary Figs. 1 – 2

Supplementary data 1 – 6 legends

**Supplementary Table 1. Summary of *Argynnia* record.** *Argynnia* species reported in the present study from Brazil, including shell length, sampling site, coordinates, biome, and type of sampled material corresponding to each specimen shown in Fig. 1.

**Supplementary Table 2. Phylogenomic matrix data information. A.** Phylogenomic Matrix (Fig. 3) Taxon Composition and Completeness Metrics. **B.** Phylogenomic Matrix (Fig. 3) Gene Composition and Indices for Concatenated Matrix. **C**. Phylogenomic Matrix (Fig. 3) Gene Occupancy Data for Each Taxon.

**Supplementary Table 3.** Blast similarity search result based on the SSU sequence of the P10K941 isolate.

**Supplementary Table 4.** Blast similarity search result based on the COI sequence of the P10K941 isolate.

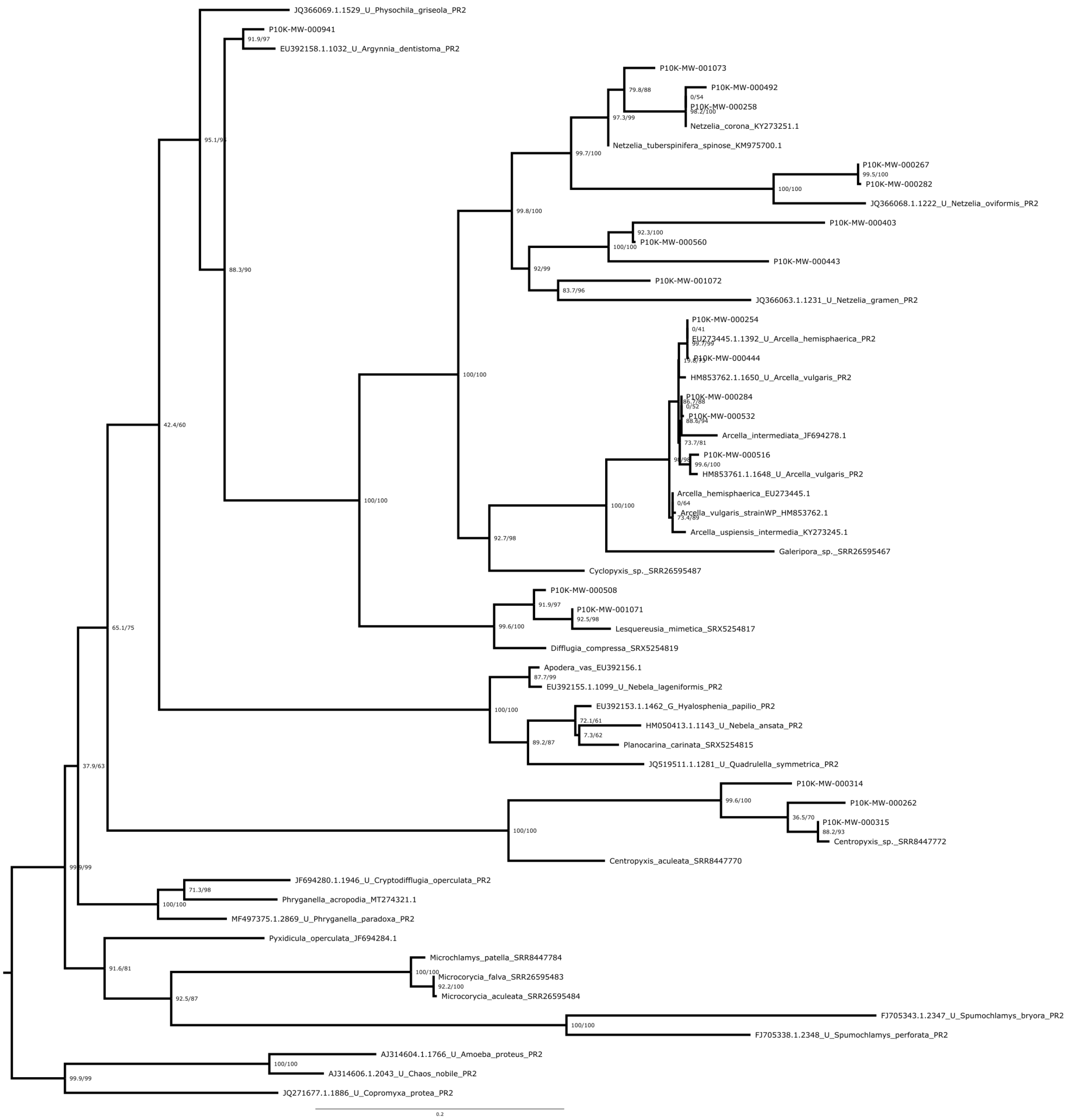


**Supplementary Fig. 1.** The maximum-likelihood phylogenetic tree of Small Subunit ribosomal RNA (SSU) representing a broad sampling of Arcellinida diversity, with three Euamoebida taxa included as outgroup. Phylogenetic reconstruction was conducted using IQ-TREE v2.3.6, with ModelFinder identifying the best-fit substitution model (TIMe+I+G4). Node support was assessed using both ultrafast bootstrap (UFBoot) and the Shimodaira–Hasegawa approximate likelihood ratio test (SH-aLRT). Support values are reported as SH-aLRT / UFBoot, with values ≥80/95 considered indicative of strong support.

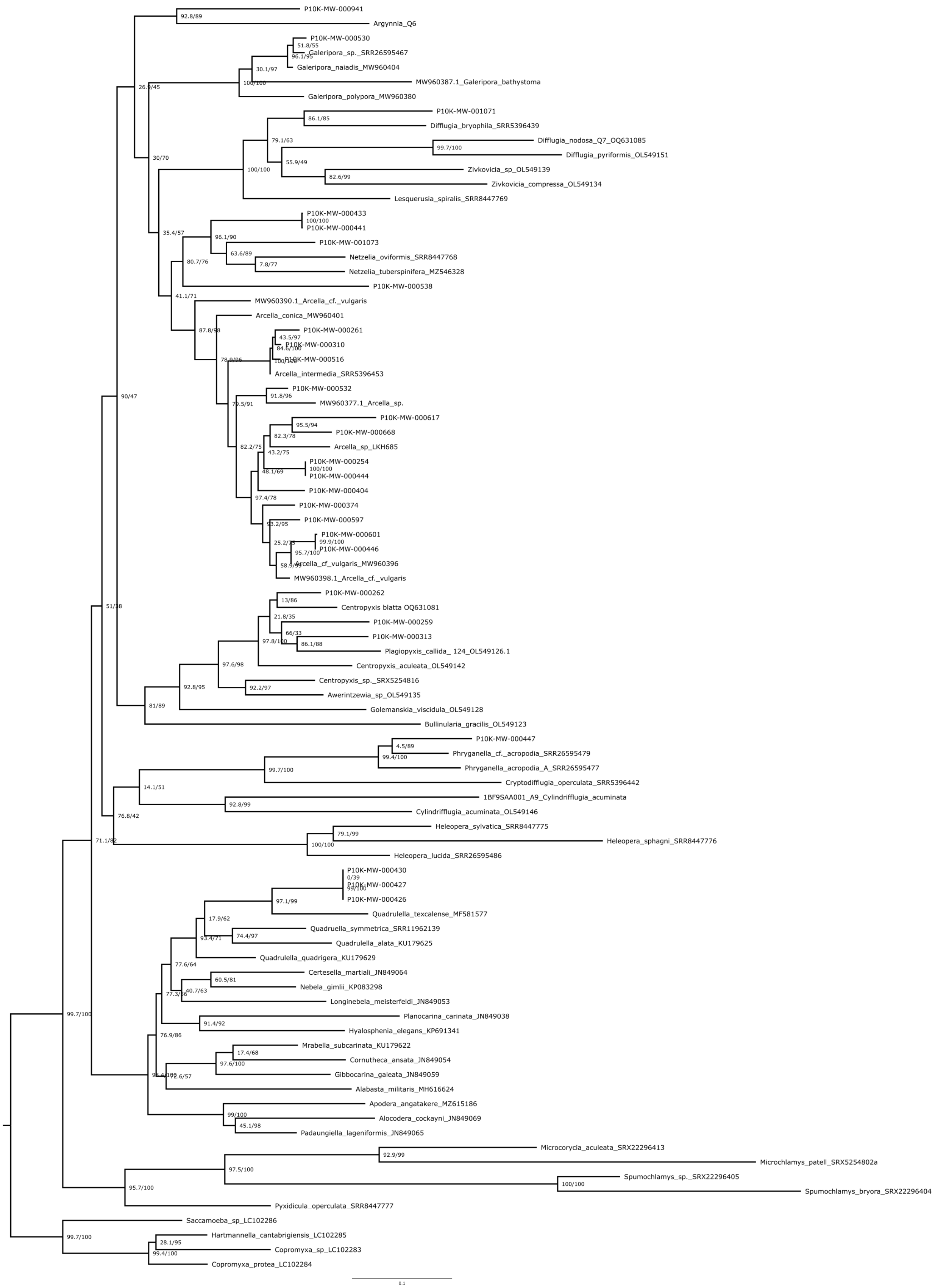

P10K-MW-000941
92.8/89
Argynnia_Q6
26.9/45
P10K-MW-000530
51.8/55
Galeripora_sp._SRR26595467
Galeripora_naiadis_MW960404
30.1/97
100/100
MW960387.1_Galeripora_bathystoma
Galeripora_polypora_MW960380
P10K-MW-001071
86.1/85
Difflugia_bryophila_SRR5396439
79.1/63
Difflugia_nodosa_Q7_OQ631085
99.7/100
Difflugia_pyriformis_OL549151
30/70
55.9/49
Zivkovicia_sp_OL549139
82.6/99
Zivkovicia_compressa_OL549134
100/100
Lesquerusia_spiralis_SRR8447769
P10K-MW-000433
100/100
P10K-MW-000441
35.4/57
96.1/90
P10K-MW-001073
63.6/89
Netzelia_oviformis_SRR8447768
7.8/77
Netzelia_tuberspinifera_MZ546328
80.7/76
P10K-MW-000538
41.1/71
MW960390.1_Arcella_cf._vulgaris
Arcella_conica_MW960401
87.8/98
P10K-MW-000261
43.5/97
P10K-MW-000310
84.6/100
P10K-MW-000516
Arcella_intermedia_SRR5396453
P10K-MW-000532
91.8/96
MW960377.1_Arcella_sp.
90/47
P10K-MW-000617
95.5/94
P10K-MW-000668
82.3/78
Arcella_sp_LKH685
82.2/75
43.2/75
P10K-MW-000254
100/100
P10K-MW-000444
48.1/69
P10K-MW-000404
97.4/78
P10K-MW-000374
P10K-MW-000597
P10K-MW-000601
99.9/100
P10K-MW-000446
95.7/100
Arcella_cf_vulgaris_MW960396
MW960398.1_Arcella_cf._vulgaris
P10K-MW-000262
13/86
Centropyxis blatta OQ631081
51/38
21.8/35
P10K-MW-000259
66/33
P10K-MW-000313
86.1/88
97.8/100
Plagiopyxis_callida_ 124_OL549126.1
97.6/98
Centropyxis_aculeata_OL549142
Centropyxis_sp._SRX5254816
92.8/95
92.2/97
Awerintzewia_sp_OL549135
81/89
Golemanskia_viscidula_OL549128
Bullinularia_gracilis_OL549123
P10K-MW-000447
4.5/89
Phryganella_cf._acropodia_SRR26595479
99.4/100
99.7/100
Phryganella_acropodia_A_SRR26595477
Cryptodifflugia_operculata_SRR5396442
14.1/51
1BF9SAA001_A9_Cylindrifflugia_acuminata
92.8/99
76.8/42
Cylindrifflugia_acuminata_OL549146
Heleopera_sylvatica_SRR8447775
79.1/99
71.1/82
100/100
Heleopera_sphagni_SRR8447776
Heleopera_lucida_SRR26595486
P10K-MW-000430
0/39
P10K-MW-000427
99/100
P10K-MW-000426
97.1/99
Quadrulella_texcalense_MF581577
17.9/62
Quadruella_symmetrica_SRR11962139
74.4/97
Quadrulella_alata_KU179625
Quadrulella_quadrigera_KU179629
77.6/64
Certesella_martiali_JN849064
60.5/81
Nebela_gimlii_KP083298
40.7/63
Longinebela_meisterfeldi_JN849053
Planocarina_carinata_JN849038
91.4/92
Hyalosphenia_elegans_KP691341
76.9/86
99.7/100
Mrabella_subcarinata_KU179622
17.4/68
Cornutheca_ansata_JN849054
97.6/100
Gibbocarina_galeata_JN849059
Alabasta_militaris_MH616624
Apodera_angatakere_MZ615186
99/100
Alocodera_cockayni_JN849069
45.1/98
Padaungiella_lageniformis_JN849065
Microcorycia_aculeata_SRX22296413
92.9/99
97.5/100
Microchlamys_patell_SRX5254802a
95.7/100
Spumochlamys_sp._SRX22296405
100/100
Spumochlamys_bryora_SRX22296404
Pyxidicula_operculata_SRR8447777
Saccamoeba_sp_LC102286
99.7/100
Hartmannella_cantabrigiensis_LC102285
28.1/95
Copromyxa_sp_LC102283
99.4/100
Copromyxa_protea_LC102284
0.1

**Supplementary Fig. 2.** The maximum-likelihood phylogenetic tree of cytochrome c oxidase subunit I (COI) representing a broad sampling of Arcellinida diversity, with four Euamoebida taxa included as outgroup. Phylogenetic reconstruction was conducted using IQ-TREE v2.3.6, with ModelFinder identifying the best-fit substitution model (GTR+F+I+G4). Node support was assessed using both ultrafast bootstrap (UFBoot) and the Shimodaira–Hasegawa approximate likelihood ratio test (SH-aLRT). Support values are reported as SH-aLRT / UFBoot, with values ≥80/95 considered indicative of strong support.

**Supplementary data legends**

**Supplementary data 1.** Small Subunit ribosomal RNA (SSU) dataset, sequences ID and NCBI accession number.

**Supplementary data 2.** Cytochrome C oxidase subunit I (COI), sequences ID and NCBI accession number.

**Supplementary data 3.** Single-protein untrimmed alignments of the 227 homologous genes considered, generated through the PhyloFisher workflow and used to construct the Tubulinea phylogenomic supermatrix (Supplementary data 6). Taxon and gene information are provided in Supplementary Table 2 of the article.

**Supplementary data 4.** Single-protein trimmed alignments of the 227 homologous genes considered, generated through the PhyloFisher workflow and used to construct the Tubulinea phylogenomic supermatrix (Supplementary data 6). Taxon and gene information are provided in Supplementary Table 2 of the article.

**Supplementary data 5.** Single-protein trees built from the trimmed alignments (Supplementary data 4) of the 227 homologous genes considered, generated through the PhyloFisher workflow. Taxon and gene information are provided in Supplementary Table 2 of the article.

**Supplementary data 6**. Tubulinea phylogenomic supermatrix composed of 227 genes (68,836 amino acid sites), constructed with the PhyloFisher workflow, and used to infer the phylogenomic tree shown in Fig. 3.
